# Deletion of the Clock Gene *Period2 (Per2)* in Glial Cells Alters Mood-Related Behavior in Mice

**DOI:** 10.1101/2020.12.09.417162

**Authors:** Tomaz Martini, Jürgen A. Ripperger, Jimmy Stalin, Andrej Kores, Michael Stumpe, Urs Albrecht

## Abstract

The circadian clock regulates many biochemical and physiological pathways, and lack of clock genes, such as *Period* (*Per*) 2, do not only affect circadian activity rhythms, but can also modulate food-anticipatory and mood-related behaviors. However, it is not known how cell-type specific expression of *Per2* contributes to these behaviors. In this study, we find that *Per2* in glial cells is important for balancing mood-related behaviors. Genetic and adeno-associated virus-mediated deletion of *Per2* in glial cells of mice leads to a depression-resistant phenotype, as manifested in reduced despair and anxiety. This is paralleled by an increase of the *GABA transporter 3* (*Gat3*) mRNA and a reduction of glutamate levels in the nucleus accumbens (NAc). Exclusive deletion of *Per2* in glia of the NAc reduced despair, but had no influence on anxiety. Our data provide strong evidence for an important role of glial *Per2* in regulating mood-related behavior.

## Introduction

Most organisms from cyanobacteria to humans have time-keeping mechanisms, termed circadian clocks, which allow adaptation to the 24-hour day (Rosbash, 2009). At the heart of this regulation lies a transcriptional-translational feedback loop, called the molecular clock. This clock is made up of a set of clock genes. In mammals, the positive arm of the clock mechanism is driven by the protein heterodimer of BMAL and CLOCK. This complex activates transcription of negative elements whose proteins Period (PER) and Cryptochrome (CRY) inhibit their own transcription by inactivating the BMAL/CLOCK transcriptional complex, thereby establishing an autoregulatory feedback loop (Takahashi, 2017). The individual cellular clocks are orchestrated in an intricate manner to establish coherent systemic rhythms (Dibner et al., 2010) that are able to provide output signals to regulate various aspects of physiology and behavior. Some of these outputs regulate mood-related behavior through modulation of neurotransmitter synthesis, uptake and degradation (Chung et al., 2014; Hampp et al., 2008; Spanagel et al., 2005), and regulation of glucocorticoid signaling (Pariante and Lightman, 2008). Absence or mutation of various clock genes have been associated with mood-related behaviors in mice and humans (Chung et al., 2014; Hampp et al., 2008; McClung et al., 2005; Partonen et al., 2007). In particular, a whole-body mutation of the clock gene *Per2* revealed a depression-resistant-like phenotype in the Porsolt’s forced swim test (FST), which was associated with reduced dopamine degradation, leading to increased dopamine levels in the nucleus accumbens (NAc) (Hampp et al., 2008). This is consistent with the view that the pharmacological manipulation of the monoaminergic system regulates mood, explaining in part the pathophysiology of depression (Nestler and Carlezon, 2006). However, current treatments for depression are often inefficient and require weeks of medication before the benefits of treatment can be observed (Cipriani et al., 2018). Since major depression is a leading cause of disability in the western world and one of the main causes of death in adolescents, with a total of 800’000 suicides due to depression annually (Disease et al., 2018), new approaches of tackling this debilitating condition are needed. An emerging new target for mood interventions are astrocytes (Zhou et al., 2019), which are in close metabolic and signaling interplay with neurons (Albrecht and Ripperger, 2018; Magistretti, 2006). In order to ensure this interplay, the two cell populations need to be precisely synchronized to each other. Systemic and cellular synchronization is one of the main tasks of the circadian clock (Dibner et al., 2010) and therefore it is not surprising that astrocytes regulate rhythmic behaviors involving clock genes (Barca-Mayo et al., 2017; Brancaccio et al., 2019; Jackson et al., 2020; Tso et al., 2017). However, whether astrocytes that lack clock genes affect mood-related behaviors is not known. Since whole-body *Per2* mutant mice display changes in the reward system (Abarca et al., 2002; Spanagel et al., 2005) and despair perception (Hampp et al., 2008), we tested mice lacking the *Per2* gene in glial cells, including astrocytes.

To this end, we generated mice lacking *Per2* in glial fibrillary acidic protein (GFAP)-positive cells by cross-breeding *Per2* floxed mice with a *Cre* mouse line. In a second approach, we deleted *Per2* in GFAP positive cells of adult animals using an adeno-associated virus (AAV) delivery approach to exclude developmental contributions in our experiments due to total lack of the *Per2* gene. Both models lacking *Per2* in *Gfap*-expressing cells (termed *GPer2* and vG*Per2* knock-out) were assessed for despair- and anxiety-related behavior using the FST and O-maze test, respectively. We found that mice from both models display a manic-like phenotype and are less anxious compared to control animals. In contrast to the whole-body *Per2* mutation, which displays a manic phenotype with reduced *monoamine oxidase A* (*Maoa*) and elevated dopamine levels (Hampp et al., 2008), the same genes associated with the monoaminergic system were normal in the *GPer2* knock-out animals. However, we observed changes in the glutamatergic and GABAergic systems.

## Results

### Deletion of *Per2* in glial cells by cross-breeding of mice

In order to study the importance of the *Per2* gene in glial cells, we crossed our floxed *Per2* (*Per2* fl/fl) animals (Chavan et al., 2016) with mice expressing the *Cre*-recombinase under transcriptional control of the human *Gfap* promoter (*GCre*) (Zhuo et al., 2001). In families where the *Cre* was inherited from the maternal side, 58% of *Cre*-negative offspring (out of 80 progeny) showed germline recombination leading to total body *Per2* heterozygous deletion. In comparison, 0% of paternally inherited *Cre* (out of 260 progeny; Supplemental table 1) showed any germline recombination, as previously described (Luo et al., 2020). Therefore, we used only male *Cre*-positive mice for matings with female *Per2* ^*fl/fl*^ animals in order to obtain glial-specific deletion of *Per2* (*GCre*^*+*^ *Per2*^*fl/fl*^ termed G*Per2*).

Next, we verified glial deletion of PER2 in brain tissue collected at zeitgeber time (ZT) 12 (where ZT0 is lights on and ZT12 is lights off) of G*Per2* mice using immunohistochemistry. Sections of the dorsal part of the suprachiasmatic nuclei (SCN) containing mainly arginine-vasopressin (AVP) neurons displayed immunoreactivity with a PER2 antibody (green) (Fig. 1A). In control animals co-staining with a GFAP antibody (red) showed partially overlapping signal with PER2 leading to yellow and orange staining, indicating PER2 expression in glial cells (white arrows, Fig. 1A, left panel). In G*Per2* mice, the red signal obtained with antibodies against the neuronal marker NeuN almost entirely overlapped with the green PER2 signal, resulting in the yellow/orange color (Fig.1A, middle panel). This indicated that in the G*Per2* animals PER2 is still present in neurons. The PER2 (green) and GFAP (red) signal did not overlap in the G*Per2* animals and no yellow color was observed (Fig. 1A, right panel), strongly suggesting that PER2 was absent in glial cells. Hence, the green signal is due to neuronal PER2 expression, suggesting specificity of our approach.

**Figure 1:**
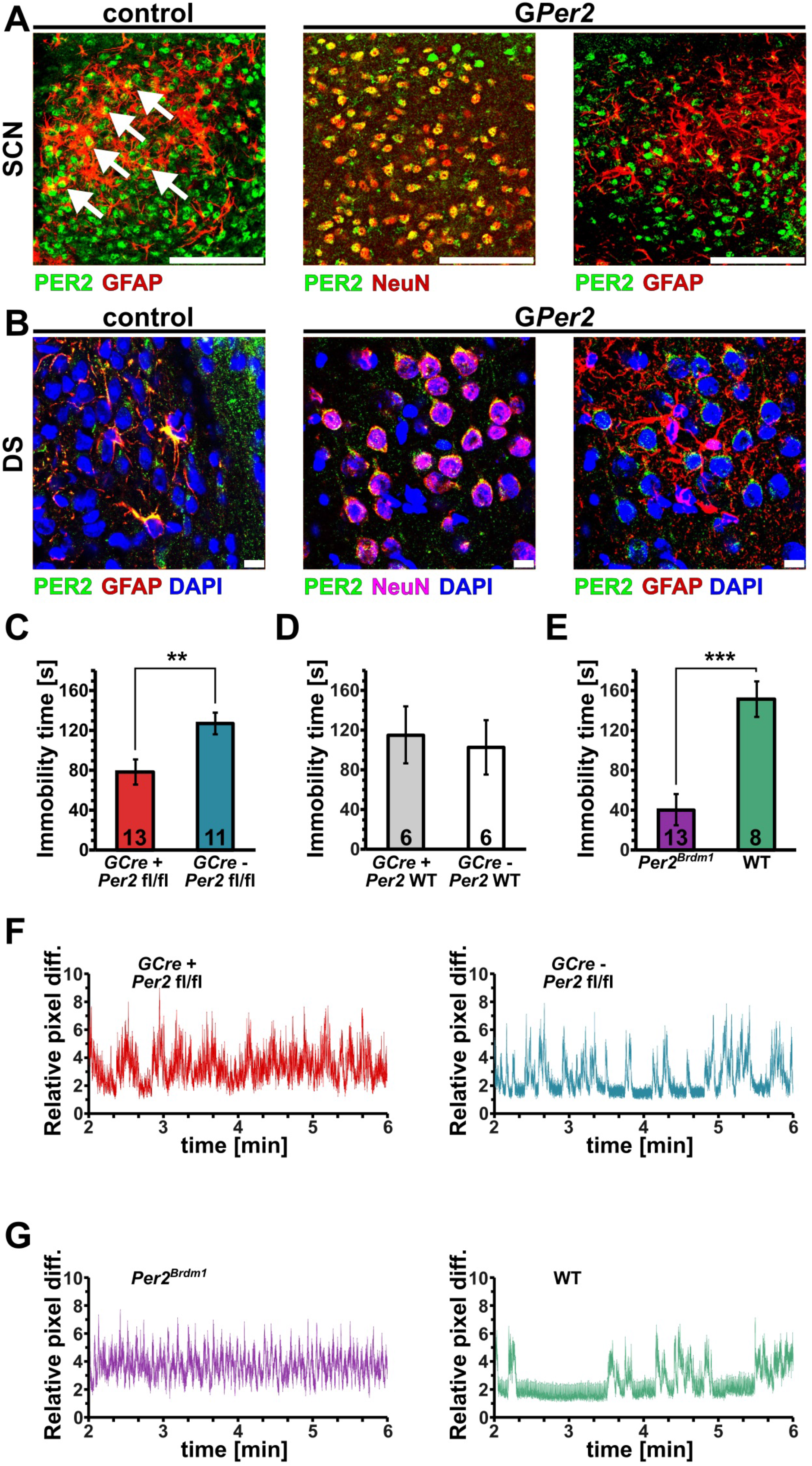
Genetic deletion of *Per2* in glia leads to depression resistant behavior. (A) Immunohistochemistry of sections from suprachiasmatic nuclei (SCN) of control (left panel) and G*Per2* mice (middle and right panels) collected at ZT12. Sections were incubated with antibodies against PER2 (green) and glial fibrillary acidic protein (GFAP), a glial marker (red, left and right panels), or NeuN (red, middle panel), a neuronal marker. White arrows in the control panel (left) indicate overlapping PER2 and GFAP signal (yellow) in glial cells, green indicates neuronal PER2 signal and red astrocytic GFAP. In G*Per2* sections, PER2 is mostly detected in neurons that are NeuN-positive, giving rise to the yellow color. In contrast, PER2 (green) is not seen in glial cells (red) of G*Per2* mice, indicating a glial specific deletion of PER2. Scale bar: 100 µm. (B) Immunohistochemistry of sections from the dorsal striatum (DS) of control (left panel) and G*Per2* mice (middle and right panels) collected at ZT12. Sections were stained with DAPI (blue) to reveal cell nuclei and were incubated with antibodies against PER2 (green) and GFAP (red, left and right panels) or NeuN (magenta, middle panel). The yellow color indicates overlapping signal of PER2 and GFAP in control animals (left panel) or PER2 and NeuN in neurons (middle panel). No yellow color in G*Per2* sections stained for PER2 and GFAP is observed. Scale bar: 10 µm. (C) Immobility time in the forced swim test (FST) of G*Per2* (*GCre+ Per2*^*fl/fl*^, red) and control (*GCre-Per2* ^*fl/fl*^, blue) animals are shown (n = 13 or 11, respectively, two-tailed t-test, **p < 0.01). (D) FST of the *Cre*-driver line (*GCre+ Per2 WT*) and the *Per2* control line (*GCre-Per2 WT*) are shown (n = 6, two-tailed t-test, no significant difference is observed). (E) FST of *Per2* mutant (*Per2*^*Brdm1*^, purple) mice and their littermate controls (WT, green) show a significant difference as previously observed (Hampp et al., 2008); n = 13 and 8, respectively, two-tailed t-test, ***p < 0.001). (F) Representative swimograms of G*Per2* (*GCre+ Per2* ^*fl/fl*^, red) and control (*GCre-Per2* ^*fl/fl*^, blue) animals. Note the longer stretches of immobility in the control animals (right panel, blue). (G) Representative swimograms of *Per2* mutant (*Per2*^*Brdm1*^, purple) mice and their littermate controls (WT, green). Note the longer stretches of immobility in the control animals (right panel, green).

To corroborate these observations, we performed immunohistochemistry in the striatum at ZT12 (Fig. 1B). The signals for PER2 (green) and GFAP (red) overlapped in the cytoplasm (yellow) and not in the nucleus (blue) of control animals (Fig. 1B, left panel). In G*Per2* animals, NeuN signal (magenta) was mainly observed around the nucleus (blue) with overlapping signal from the PER2 antibody (green) resulting in the yellow color (Fig. 1B, middle). Note that there was no green signal observed that could stem from glial PER2 expression. In the G*Per2* mice the PER2 signal (green) did not overlap with the GFAP signal (red) and PER2 appeared to be mainly present around the nuclei (blue) of neurons. Overall, our immunohistochemistry data indicate that PER2 was absent from glial cells of G*Per2* mice.

### G*Per2* mice display a depression-resistant-like phenotype

The glial *Per2* knock-out animals (G*Per2*) were tested for despair-based behavior, which is one of the important manifestations of depression. To this end, we used the forced swim test (FST) to assess time of immobility at ZT6 (Hampp et al., 2008). We observed that G*Per2* mice (*GCre*^*+*^ *Per2*^*fl/fl*^) were significantly less immobile compared to control animals (*GCre*^*-*^ *Per2*^*fl/fl*^) (Fig. 1C). The *Cre*-driver animals used for crossing (*GCre*^*+*^ *Per2*^*WT*^ and *GCre*^*-*^ *Per2*^*WT*^) showed an immobility time comparable to the control animals (*GCre*^*-*^ *Per2*^*fl/fl*^) (Fig. 1D). This illustrated that the lower immobility observed in the G*Per2* mice was specific to the lack of *Per2* in glial cells. As a positive control, we reproduced the lower immobility phenotype of *Per2*^*Brdm1*^ mutant mice (Fig. 1E) as described previously in male mice (Hampp et al., 2008), but this time using female animals to demonstrate that the phenotype was not gender specific. Figure 1F shows examples of primary data depicted as mobility over time (swimograms) with the corresponding color-coding. The data were obtained using a self-developed movement analysis software to determine mobility in an automated unbiased manner (see methods). It was evident that the control animals displayed long stretches of continuous baseline signal, which corresponded to long time stretches of immobility (Fig. 1F, G, right panels). These long stretches of immobility were very short or absent in both the G*Per2* (*GCre*^*+*^ *Per2*^*fl/fl*^) and *Per2*^*Brdm1*^ mutant mice (Fig. 1F, G, left panels). Taken together, our data show that lack of *Per2* in glial cells was sufficient to reproduce the manic-like phenotype observed previously in whole-body *Per2* mutant mice (Hampp et al., 2008). This suggests that *Per2* in glial cells may play an important role in the development of despair-based behavior contributing to depression.

### Deletion of *Per2* in glial cells using adeno-associated virus (AAV)-delivered *Cre* recombinase

The *Gfap*-Cre driver mouse line used above has been reported to potentially express *Cre* in some neuronal progenitor cells (Harno et al., 2013; Quintana et al., 2012). Furthermore, the *Cre* recombinase is most often inserted randomly into the genome and its correct position and number of copies are unknown. Additionally, *Cre* leakage was observed in some cases leading to unwanted recombination events (Lindhorst et al., 2020; Song and Palmiter, 2018). Due to these potential problems and to exclude developmental effects in our deletion approach above, we delivered a construct expressing the codon-improved *Cre*-recombinase (*iCre*) using engineered adeno-associated viruses (AAVs). An engineered AAV for efficient non-invasive gene delivery that can cross the blood-brain barrier (BBB) was used (Chan et al., 2017).

This AAV-PHP.eB contained a vector expressing *iCre* under control of the human *Gfap* promoter with an enhanced green fluorescent protein (*eGfp*) as reporter. After intravenous (i.v.) injection via the lateral tail vein, fluorescence in the brain was detectable after 3 weeks in both the synthetic CAG-driven positive control (Miyazaki et al., 1989), as well as in the *Gfap*-driven constructs (Fig. 2A, red and yellow signals). No fluorescence and hence no BBB permeability was detected when using the natural variant AAV9 to deliver the CAG-driven control. Similarly, the non-injected control brain showed only baseline signal (Fig. 2A, brown). Intraperitoneal (i.p.) injection of the AAV-PHP.eB *Gfap*-driven *Cre*-construct also showed only baseline signal after 3 weeks. After 2 months, however, even i.p. delivery of the AAV-PHP.eB *Gfap*-driven construct resulted in detectable fluorescene, comparable to that of the i.v. injection (Fig. 2B, right), which is quite remarkable and is shown here to work for the first time. We termed the AAV mediated deletion of *Per2* in glial cells vG*Per2*.

**Figure 2.**
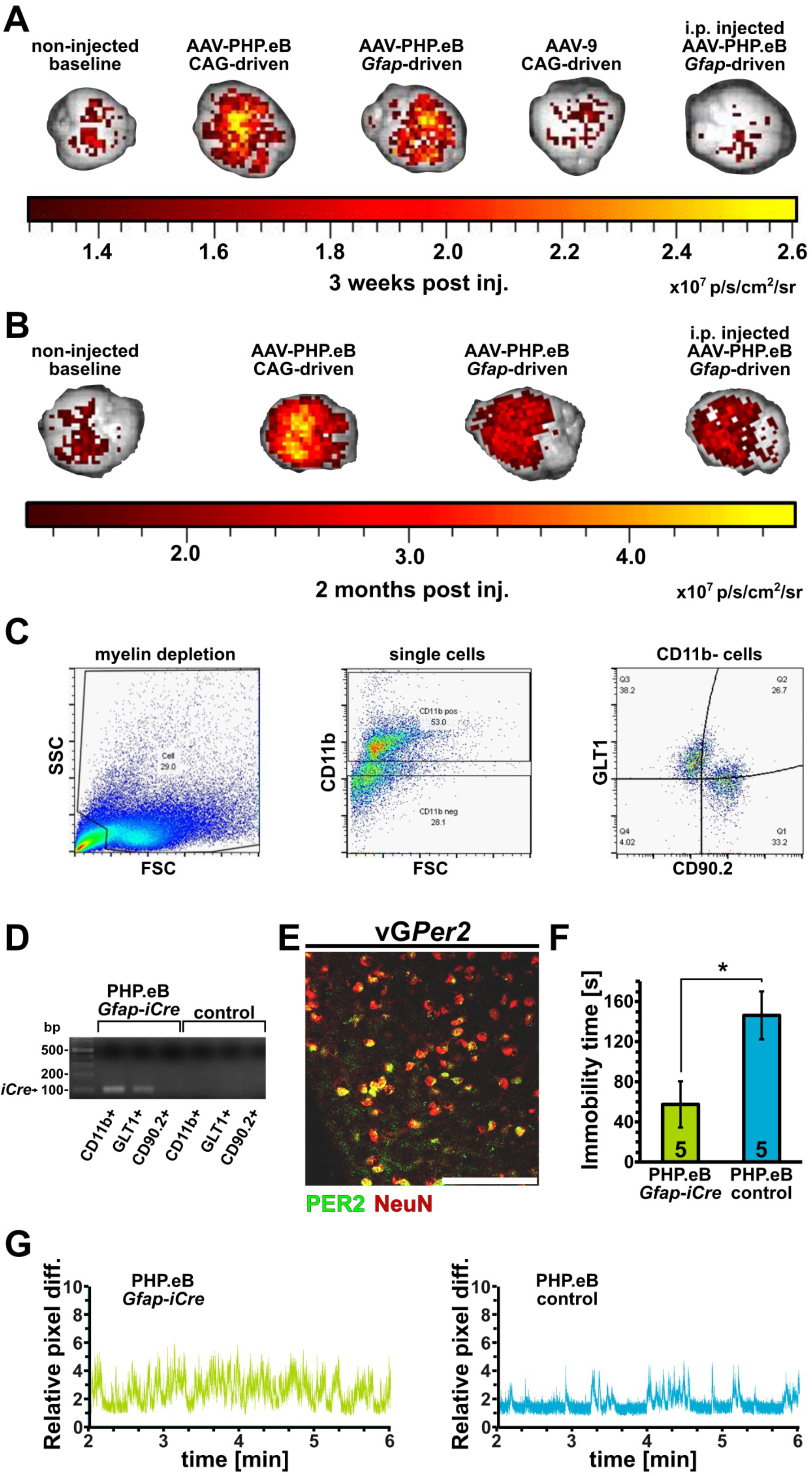
Adeno-associated virus (AAV)-mediated deletion of *Per2* in glial cells of adult mice leads to a depression resistant behavior. (A) Fluorescent imaging of whole brains 3 weeks after no injection (left), intravenous (i.v.) injection of the engineered AAV-PHP.eB, which can pass the blood-brain barrier (BBB), containing the general *CAG* driver (second from left) or the glial *Gfap* driver (middle). The second to last brain is from an animal with i.v. injected AAV-9, which does not pass the BBB, containing the general *CAG* driver. The last brain (right) is from an animal injected intraperitoneally (i.p.) with the AAV-PHP.eB *Gfap*-driven construct. Note that only the brains of animals that received the AAV-PHP.eb i.v. display significant fluorescent signal after 3 weeks (orange and yellow color). (B) Fluorescent imaging of whole brains 2 months after injection of the AAV-PHP.eB. Note that the fluorescence is still maintained after 2 months post injection and that even the i.p. injected AAV-PHP.eB *Gfap* is showing signal in the brain now. (C) Sorting of neurons and astrocytes by flow cytometry from brain tissue including the nucleus accumbens (NAc). The left panel shows the removal of debris from a single cell suspension, showing the distribution of debris in the forward as well as in the side scatter (FSC and SSC, respectively). The middle panel shows the removal of CD11b^+^ cells (microglia) from the cell suspension (lower left corner from left panel). The CD11b^-^ cells (bottom half from middle panel) were then sorted into two distinct cell populations corresponding to astrocytes (GLT1^+^/CD90.2^-^) and neurons (CD90.2^+^/GLT1^-^) (right panel). (D) PCR analysis of astrocytes and neurons from the cell sorting. Microglia (CD11^+^) as well as astrocytes (GLT1^+^/CD90.2^-^), but not neurons (CD90.2^+^/GLT1^-^) from PHP.eB *Gfap-iCre* infected animals show the presence of *iCre*, indicating that only glia and not neurons could express *iCre* in order to delete *Per2* in the *Per2* ^*fl/fl*^ mice. (E) Immunohistochemistry of vG*Per2* brain tissue from nucleus accumbens (NAc) isolated at ZT6. The signal for PER2 (green) mainly overlaps with neuronal NeuN signal (red) giving rise to the yellow color. Scale bar: 100 µm. (F) Immobility time in the forced swim test (FST) of vG*Per2* (PHP.eB *Gfap-iCre*, green) and control (PHP.eB control, blue) animals are shown (n = 5, two-tailed t-test, *p < 0.05). (G) Representative swimograms of vG*Per2* (AAV-PHP.eB *Gfap-iCre*, green) and control (AAV-PHP.eB control, blue) animals. Note the longer stretches of immobility in the control animals (right panel, blue).

Next, we wanted to check if the approach was specific for glial cells (astrocytes and microglia). We used flow cytometry and fluorescence-activated cell sorting to separate microglia (CD11b^+^), astrocytes (GLT1^+^) and neurons (CD90.2^+^). In a first step, the cells were depleted from debris (Fig. 2C, left panel), then the CD11b^+^ microglia were removed (Fig. 2C, middle panel) and subsequently the GLT1^+^ astrocytes were separated from the CD90.2^+^ neurons (Fig. 2C, right panel). The presence or absence of *iCre* was then determined by PCR on the different cell populations. We observed that the PHP.eB *Gfap*-*iCre* infected microglia (CD11b^+^) and astrocytes (GLT1^+^) expressed *iCre*. In contrast, neurons (CD90.2^+^) and non-infected controls did not express *iCre* (Fig. 2D). Unfortunately, the sensitivity was not sufficient to reliably detect *Per2* expression in WT non-infected controls using this approach. However, immunohistochemistry on nucleus accumbens tissue isolated at ZT6 confirmed that PER2 protein could still be detected in neurons, while outside of neurons PER2 was barely or not detected (Fig. 2E).

### vG*Per2* mice display a depression-resistant-like phenotype

The animals with intravenously (i.v.) applied AAV-PHP.eB, which mediated the deletion of *Per2* in glial cells (vG*Per2*), were subjected to the FST in order to assess their immobility in this despair-based behavioral test. We observed that vG*Per2* (PHP.eB *Gfap-iCre*) mice showed significantly lower immobility times compared to control animals, which were injected with a comparable virus, but lacking *iCre* (PHP.eB control) (Fig. 2F). Examples of swimograms illustrating swimming behavior between minute 2 to 6 are shown (Fig. 2G, Suppl. Fig. 1). Longer resting bouts were only detected in control animals (Fig. 2G, right panel, Suppl. Fig. 1). This was consistent with the result we obtained by cross-breeding *Per2* floxed with *Gfap*-*Cre* mice (Fig. 1C, F, G), indicating that the phenotype can be reproduced by deleting *Per2* in glial cells of the adult animal. Hence, developmental processes are very unlikely to be responsible for this depression-resistant-like phenotype.

### G*Per2* and vG*Per2* mice show reduced anxiety-like behavior in the O-maze

Depression is a complex state and in addition to despair-related aspects also involves features of anxiety. Therefore, we tested the G*Per2* as well as the vG*Per2* animals in the elevated O-maze and measured how much time they spent in the open area of the maze and how many times they entered the open part. We observed that both the G*Per2* as well as the vG*Per2* animals spent more time in the open section of the O-maze compared to control animals (Fig. 3A, B). Interestingly, entries into the open section of the O-maze were not significantly different compared to controls (Fig. 3C, D), although both G*Per2*, as well as vG*Per2* mice, displayed a tendency to explore the open section more frequently. These observations indicate a reduced anxiety level with a tendency of higher exploratory behavior in mice that lack *Per2* in glial cells.

**Figure 3.**
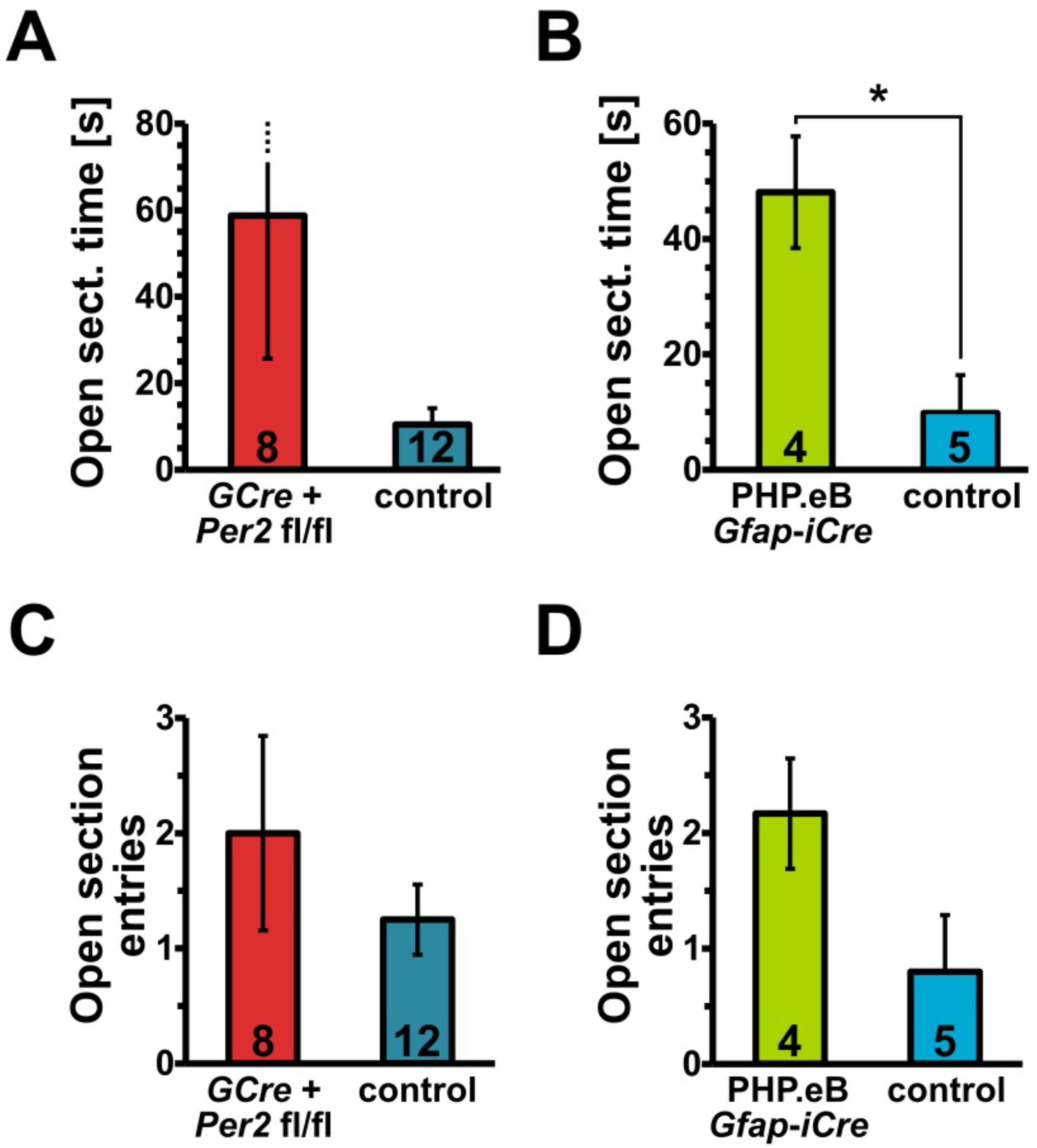
Reduced anxiety of G*Per2* and vG*Per2* animals in the elevated O-maze. (A) Time spent in the open area of the O-maze is slightly longer in G*Per2* (*GCre+ Per2 fl/fl*, red) animals compared to controls (blue), however, this difference is just a trend (n = 8 and 12, respectively, two-tailed t-test, p = 0.09). (B) Time spent in the open area of the O-maze is significantly longer in vG*Per2* (AAV-PHP.eB *Gfap-iCre*, green) compared to controls (AAV-PHP.eB control, blue) (n = 4 and 5, respectively, two-tailed t-test, *p < 0.05). (C) Entries into the open section of the O-maze are not significantly different between the G*Per2* animals (red) and their controls (blue) (n = 8 and 12, respectively, two-tailed t-test, p = 0.35). (D) Entries into the open section of the O-maze is not significantly different between the vG*Per2* animals (green) and their controls (blue) (n = 4 and 5, respectively, two-tailed t-test, p = 0.08).

### G*Per2* mice display normal circadian parameters

Since G*Per2* mice display less immobility in the FST and swim more, we tested these animals for the circadian parameters of total activity, circadian period and body temperature fluctuation. We measured running-wheel revolutions in the cages of G*Per2* and control animals and observed identical activity profiles with low activity during the 12-hour light phase and high activity during the 12-hour dark phase (Fig. 4A). General activity was measured via an intraperitoneally implanted transmitter that was traced by a detector plate under the floor of the cage. Comparable to the wheel-running activity (Fig. 4A), no difference between the genotypes was observed (Fig. 4B). However, the activity in the second half of the dark phase (ZT16-22) was for both genotypes higher in the wheel-running assessment compared to the general activity pattern. These results show, that the reduced immobility time of G*Per2* animals in the FST was not due to a higher activity level compared to controls and, hence, the higher activity in the FST is related to despair rather than to general activity.

**Figure 4.**
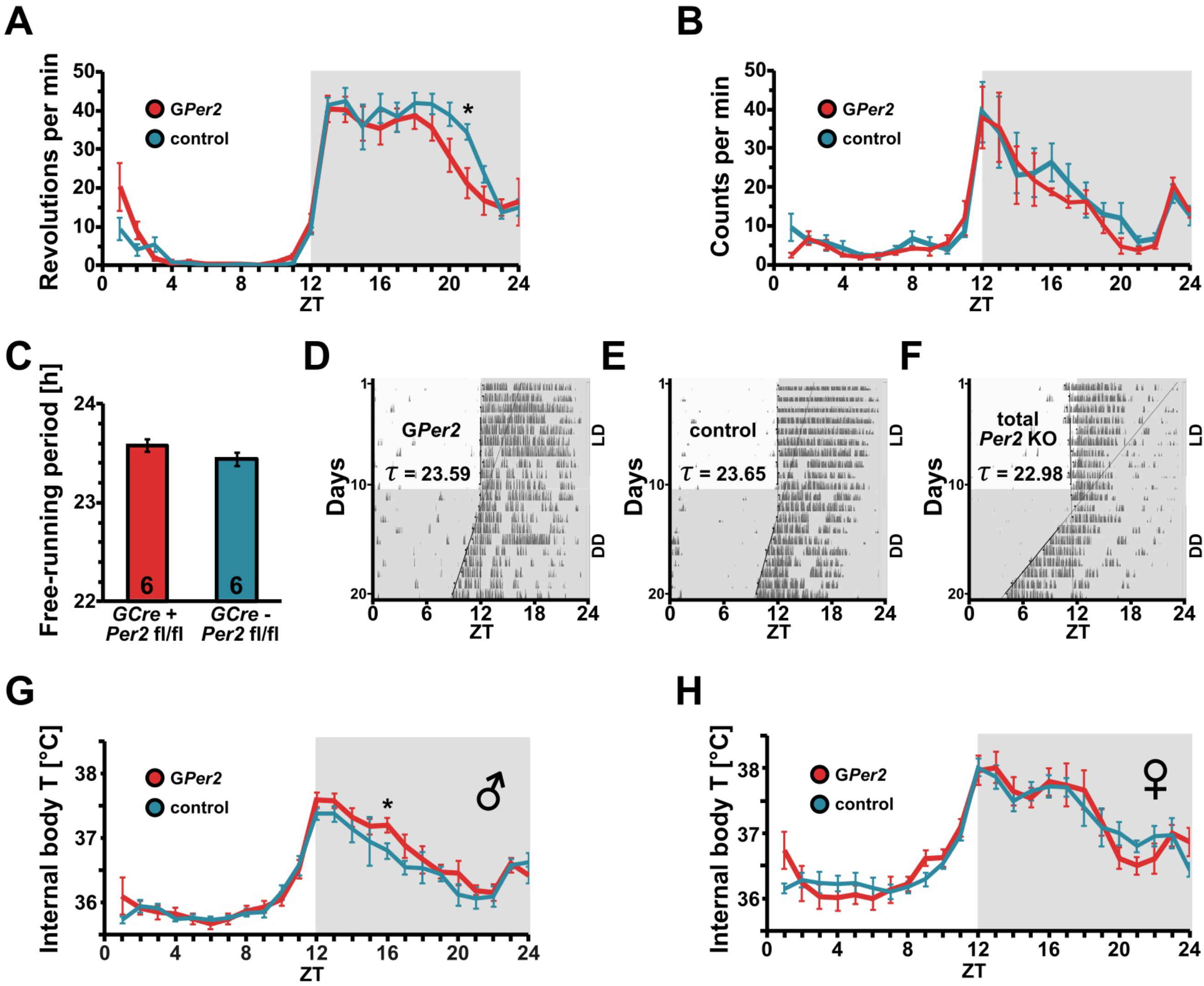
G*Per2* mice have a normal circadian clock. (A) Wheel-running activity profile of G*Per2* (*GCre+ Per2* ^*fl/fl*^, red) and control (*GCre-Per2* ^*fl/fl*^, blue) mice. The profiles are almost identical with a slight reduction of activity in G*Per2* animals at zeitgeber time (ZT) 21. (B) General activity pattern of G*Per2* (red) and control (blue) mice. The two profiles are not significantly different. (C) Circadian period of G*Per2* (red) and control (blue) mice. No significant difference was observed between the two genotypes (n = 6, two-tailed t-test, p > 0.05). (D) Representative wheel-running actogram of a G*Per2* animal. Black vertical marks represent activity in the wheel. Upper part shows the activity under an LD 12:12 cycle. Lower part shows the activity under constant darkness conditions (DD). (E) Representative wheel-running actogram of a control animal. (F) Representative wheel-running actogram of a whole-body total *Per2* knock-out animal. (G) Body temperature profile of male G*Per2* (red) and control (blue) mice under LD conditions (n = 6). (H) Body temperature profile of female G*Per2* (red) and control (blue) mice under LD conditions (n = 6).

Using the wheel-running activity data, we also determined the circadian period (τ) of the G*Per2* and total *Per2* knock-out (KO) animals under constant darkness conditions. The circadian period of G*Per2* mice was normal and not significantly different from control animals (Fig. 4C, D, E), indicating that loss of *Per2* in glial cells does not affect circadian period. In contrast, whole-body *Per2* KO mice displayed a period shorter than 23 hours (Fig. 4F), consistent with previous findings (Chavan et al., 2016; Zheng et al., 1999).

Next, we assessed body temperature over 24 hours under a 12-hour light : 12-hour dark cycle (LD 12:12). No significant differences in the body temperature profiles were observed between G*Per2* and control animals, in both males (Fig. 4G) as well as females (Fig. 4H). This suggested that loss of *Per2* in glial cells does not affect body temperature regulation.

Taken together, these data illustrate that loss of *Per2* in glial cells does not phenocopy all the characteristics of *Per2* whole-body KO animals. This highlights a specific role of glial *Per2* in mood-related behaviors.

### Molecular changes in G*Per2* mice

Mood-related behaviors including depression are regulated by a number of different brain nuclei and regions (Altshuler et al., 2010; Banasr and Duman, 2008; Bhagwagar et al., 2008; Duncan et al., 1993; Etievant et al., 2015; Zink et al., 2009). We investigated three of those brain regions, the nucleus accumbens (NAc), the medial prefrontal cortex (mPFC) and the amygdala (AMY), as well as the hypothalamus (HYP) as a control brain region. The aforementioned brain regions also showed altered activity in response to swim stress (Duncan et al., 1993). We focused our attention on expression of genes involved in the synthesis, reuptake and degradation of monoamine neurotransmitters, as well as on genes involved in the clearance of glutamate and GABA from the synaptic cleft, because some of these processes were observed to be altered in *Per2* mutant mice (Hampp et al., 2008; Spanagel et al., 2005). Special attention was given to astrocyte-specific genes (Supplemental table 2).

In the hypothalamus (HYP), *GABA transporter 1* (*Gat1*) mRNA was significantly decreased in G*Per2* (*GCre*^*+*^ *Per2* ^*fl/fl*^) compared to control (*GCre*^*-*^ *Per2*^*fl/fl*^) animals (Fig. 5A). In the nucleus accumbens (NAc), mRNA expression of *GABA transporter 3* (*Gat3*) and *dopamine receptor D3* (*Drd3*) was increased in G*Per2* (*GCre*^*+*^ *Per2* ^*fl/fl*^) compared to control (*GCre*^*-*^ *Per2*^*fl/fl*^) mice (Fig. 5B, C). Interestingly, genes coding for enzymes involved in monoamine synthesis, such as *tyrosine hydroxylase* (*Th*) and monoamine degradation, such as *monoamine oxidases A* (*Maoa*) and *B* (*Maob*), as well as *catechol-O-methyltrasferase* (*Comt*) were similar between the two genotypes in the NAc (Fig. 5D-G), while *Per2* mRNA was significantly reduced in G*Per2* mice (Fig. 5H). These results suggest that *Per2* in glial cells is involved in the regulation of GABA signaling, rather than the regulation of monoaminergic signaling. This was further underlined by our findings on the analysis of neurotransmitters. We found that glutamate (Glu) levels were significantly decreased in the NAc of G*Per2* (*GCre*^*+*^ *Per2* ^*fl/fl*^) mice (Fig. 5I). This difference, however, was not significant in the other brain areas investigated (dorsal striatum (DS), medial prefrontal cortex (mPFC), Supplemental table 3). We did also not observe changes in GABA and glutamine in the NAc, DS and mPFC (Supplemental table 3).

**Figure 5.**
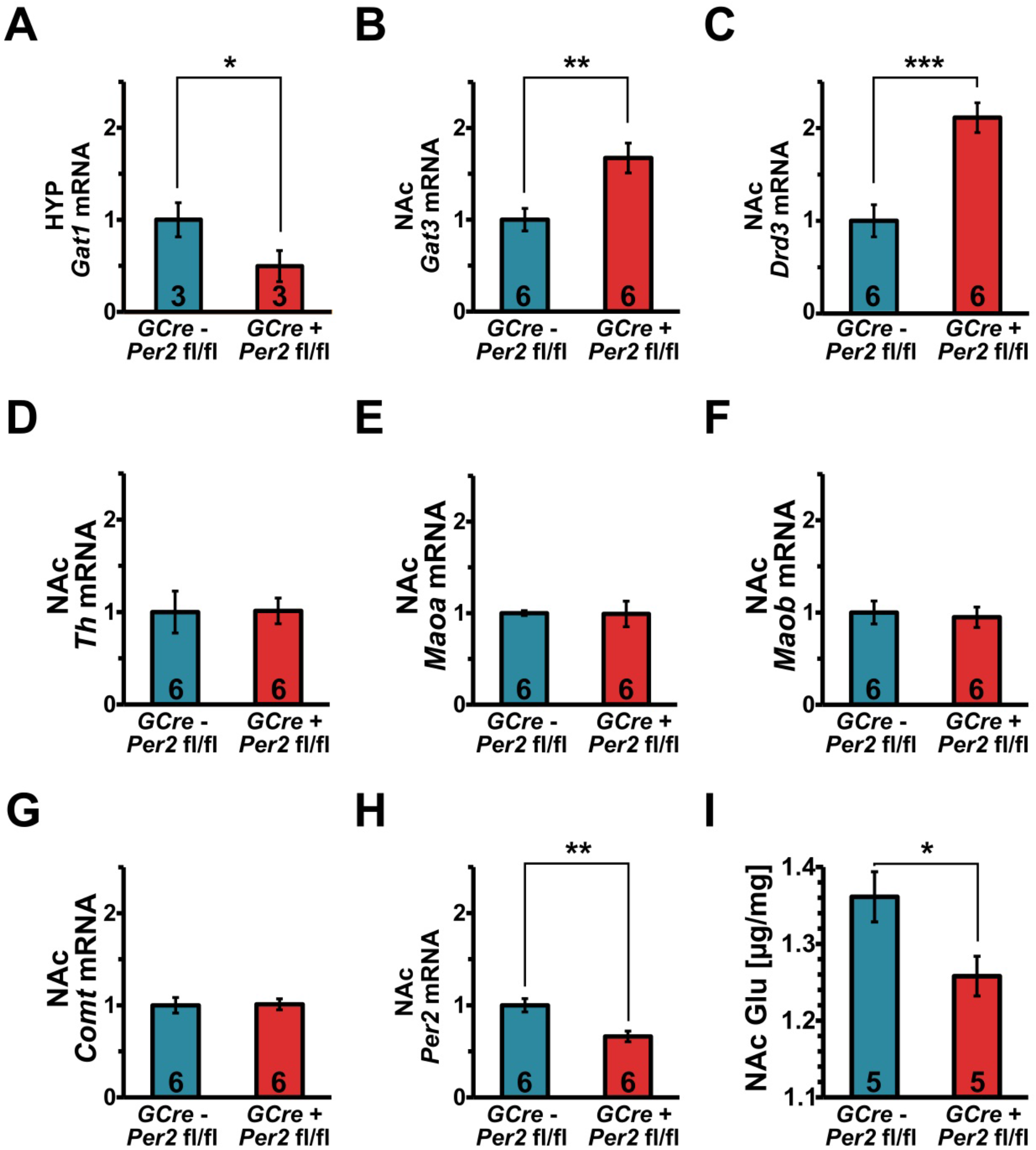
Molecular changes in G*Per2* mice. (A) Expression of *GABA transporter 1* (*Gat1*) mRNA in the hypothalamus (HYP) of G*Per2* (*GCre+ Per2* ^*fl/fl*^, red) control (*GCre-Per2* ^*fl/fl*^, blue) at ZT2 (n = 3, two-tailed t-test, *p < 0.05). (B) Expression of *GABA transporter 3* (*Gat3*) mRNA in the nucleus accumbens (NAc) of G*Per2* (red) and control (blue) animals (n = 6, two-tailed t-test, **p < 0.01). (C) Expression of *dopamine receptor D3* (*Drd3*) mRNA in the NAc of G*Per2* (red) and control (blue) animals (n = 6, two-tailed t-test, ***p < 0.001). (D) Expression of *tyrosine hydroxylase* (*Th*) mRNA in the NAc of G*Per2* (red) and control (blue) animals (n = 6, two-tailed t-test, p > 0.05). (E) Expression of *monoamine oxidase A* (*Maoa*) mRNA in the NAc of G*Per2* (red) and control (blue) animals (n = 6, two-tailed t-test, p > 0.05). (F) Expression of *Maob* mRNA in the NAc of G*Per2* (red) and control (blue) animals (n = 6, two-tailed t-test, p > 0.05). (G) Expression of *catechol-O-methyltrasferase* (*Comt*) mRNA in the NAc of G*Per2* (red) and control (blue) animals (n = 6, two-tailed t-test, p > 0.05). (H) Amount of glutamate (Glu) in the NAc of G*Per2* (red) and control (blue) animals at ZT6 (n = 5, two-tailed t-test, *p < 0.05).

Taken together, we provide evidence that lack of *Per2* in glia affects mostly the NAc, and there the signaling pathways involving GABA and glutamate.

### Deletion of *Per2* in glial cells of the NAc is sufficient to elicit a depression-resistant-like phenotype, but does not affect anxiety-like behavior

Since we observed most of the molecular and neurotransmitter changes in the NAc of G*Per2* animals, we wondered whether deletion of *Per2* in glial cells of the NAc alone could elicit the phenotypes described above. Therefore, we injected our viral vectors directly into the NAc of *Per2*^*fl/fl*^ mice (Fig. 6A) using a stereotactic injection apparatus and termed the animals GNAc*Per2* mice. We saw that both AAV vectors, the PHP.eB *Gfap-iCre* as well as the AAV9 *Gfap-iCre* significantly reduced immobility time in the FST (Fig. 6B, C). The results were comparable to the observations in G*Per2* and the vG*Per2* animals, in which *Per2* was deleted in glial cells throughout the brain (Fig. 1 and 2). In contrast, we did not observe an effect on the time spent in the open section of the O-maze when AAV9 *Gfap-iCre* was injected into the NAc (Fig. 6D), suggesting that glial *Per2* in the NAc was not involved in the regulation of anxiety-related behavior.

**Figure 6.**
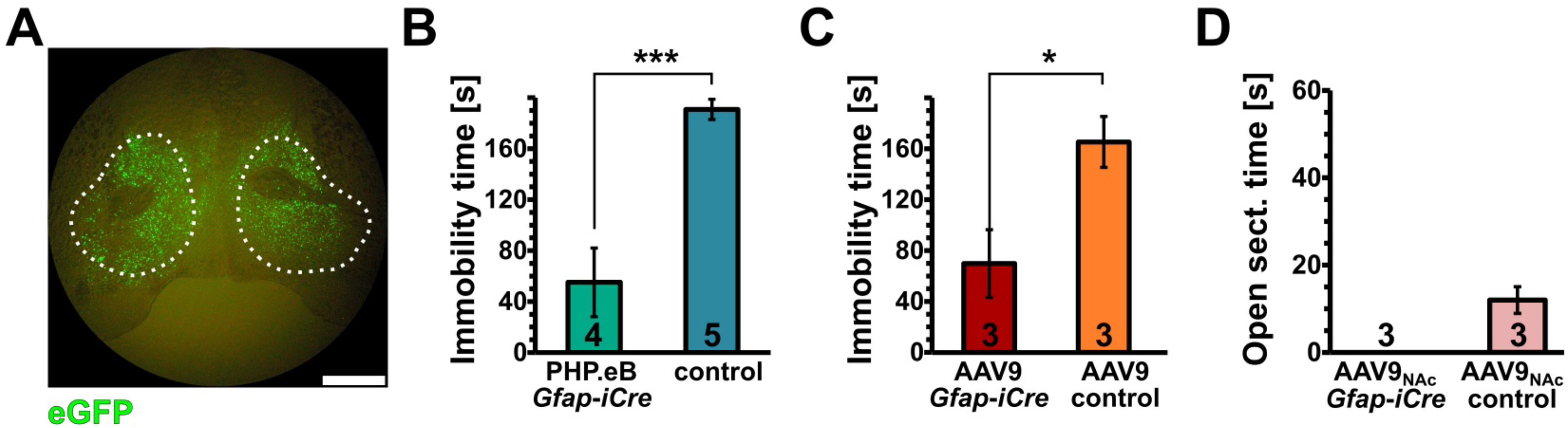
Deletion of *Per2* in glia of the NAc evokes depression resistant behavior but has no effect on anxiety-related behavior. (A) Injection of the AAVs into the NAc as revealed by the eGFP reporter fluorescence. White dotted area delineates the NAc. Scale bar: 1 mm. (B) Immobility time in the forced swim test (FST) of PHP.eB *Gfap-iCre* (green) and non-injected control (*Per2* ^*fl/fl*^, blue) animals is shown (n = 4 or 5, respectively, two-tailed t-test, ***p < 0.001). (C) FST of AAV9 *Gfap-iCre Per2* ^fl/fl^ (red) and *Gfap*-*iCre Per2* WT littermate control (orange) NAc injected mice is shown (n = 3, two-tailed t-test, *p < 0.05). Deletion of glial *Per2* in the NAc with both types of AAVs appears to be sufficient to evoke a depression-resistant behavior. (D) Time spent in the open area of the O-maze is similar in both NAc injected AAV9 *Gfap-iCre* and AAV9 control animals, suggesting no effect of *Per2* glial deletion in the NAc on anxiety-related behavior (n = 3, two-tailed t-test, p > 0.05).

Taken together, our data provide evidence that *Per2* expression in glial cells of the NAc is responsible for the regulation of despair-based behavior, but is not sufficient to modulate the anxiety aspect of depression.

## Discussion

Mood-related behaviors including depression are determined by several and sometimes synergistic factors illustrating the multifactorial nature of these neurological disturbances. In addition to the molecular complexity of these processes, cellular heterogeneity and topological organization complicate the understanding of the functioning of the brain even further. Previous studies have shown that at the molecular level genes involved in the regulation of the circadian clock can affect mood-related behaviors in mice (Chung et al., 2014; Hampp et al., 2008; McClung et al., 2005) and most likely also in humans (Li et al., 2013; Partonen et al., 2007). Anatomical and genetic tracing studies have also revealed particular brain regions to be involved in various aspects of mood regulation.

Our previous studies have implicated a role of the clock gene *Per2* in mood-related behavior involving the mesolimbic dopaminergic system (Hampp et al., 2008). However, the contribution of glial cells in this process is poorly understood (Zhou et al., 2019). Therefore, we were interested to investigate the role of glial *Per2*. We used three approaches to investigate this question. First, we used genetic tools to delete *Per2* specifically in glial cells, including astrocytes (Fig. 1). Second, we deleted *Per2* in glial cells of the adult animal via systemic application of an engineered AAV that could pass the BBB (Fig. 2). This experiment was performed to exclude potential developmental effects that could contribute to behavioral alterations in the genetic approach. Third, we deleted Per2 specifically in glial cells of the NAc by injection of AAVs expressing *Cre* recombinase. All three approaches, the genetic as well as the AAV approaches revealed that absence of *Per2* in glial cells, in particular the NAc, reduced the immobility time of mice in the FST, a test that assesses despair-based behavior, one of the aspects of depression. Glial deletion of *Per2* resulted in more active swimming in the FST, correlating with a manic behavior as we have previously observed in whole-body *Per2* mutant mice (Hampp et al., 2008). Interestingly, pharmacologic glial ablation in the prefrontal cortex has been reported to increase immobility in the FST and as a consequence made the animals more depressive (Banasr and Duman, 2008). This phenotype is the opposite from what we observed in our G*Per2* and vG*Per2* mice and indicates that in our animal models, glial cells are still functional and not eliminated. Hence, our observations are specific to glial *Per2* function and are very unlikely related to processes leading to glial cell death.

The processes affected by lack of *Per2* in glial cells are most likely not related to the circadian clock mechanism, because G*Per2* animals display no abnormalities in clock parameters such as period or activity distribution and body temperature fluctuations (Fig. 4). If the disturbance of the clock mechanism in astrocytes would play a major role, a desynchronization between astrocytes and neurons would be expected, which would ultimately lead to arrhythmic activity of mice under constant darkness conditions. This is, however, not what we observed (Fig. 4). Therefore, we hypothesize that the phenotypes we describe here are probably related to PER2 output functions, e.g. as nuclear receptor co-regulator (Schmutz et al., 2010). Potential nuclear receptor binding sites could be detected in the *Gat3* promoter using the Genomatix software suite (Cartharius et al., 2005). PER2 may potentially interact with some of these candidates and act as a nuclear receptor co-regulator, which would correlate with the altered levels of *Gat3* mRNA that we observed in the NAc (Fig. 5B). However, elaboration of this mechanism remains a topic of future investigations and may reveal novel pharmacologic targets for the treatment of mood disorders.

Since depression is not only related to despair, we also tested our animals in the elevated O-maze. This test addresses anxiety-related behavior, which is an important component of depression. We observed that both G*Per2* as well as vG*Per2* mice were less anxious, because they spent more time in the open area of the O-maze (Fig. 3). Interestingly, however, the animals with *Per2* deleted in the glia of the NAc only (GNAc*Per2*) did not show this phenotype (Fig. 6D). From this we conclude that *Per2* in glia of the NAc may not be important for anxiety-related behavior and specifically affects despair. Hence, glial *Per2* in other brain regions, such as the amygdala (Davis, 1992), the bed nucleus of the strai terminalis (BNST) (Rodriguez-Romaguera et al., 2020) or the subgenual anterior cingulate cortex (sgACC) (Alexander et al., 2020), part of the medial prefrontal cortex (mPFC), may be involved in the modulation of anxiety. We did not test anhedonia, another aspect of depression, because whole-body *Per2* mutant mice did not show any changes in the sucrose preference test (Spanagel et al., 2005). Therefore, we assumed that glial *Per2* plays a subordinate or no role in the anhedonia aspect of depression.

Our previous studies describing whole-body *Per2* mutant mice highlighted a functional involvement of *Per2* in the regulation of monoamine degradation (Hampp et al., 2008). In this study, we find that glial *Per2* appears to play a role in the regulation of glutamate and GABA signaling, but not monoamine metabolism (Fig. 5). This suggests that the role of *Per2* in neurons may relate to the dopaminergic system, whereas its function in glial cells involves glutamate and GABA signaling. There is ample evidence that both, monoaminergic and GABAergic signaling regulate depressive disorders (Luscher et al., 2011), and *Per2* might be involved in both. If this hypothesis is correct, then we would expect that deletion of *Per2* in neurons would still affect mood-related behaviors, but the signaling involved would mainly relate to the monoaminergic system, including dopamine. However, the future will reveal whether this is indeed the case.

Our finding that lack of glial *Per2* led to increased expression of the glia specific *GABA transporter 3* (*Gat3*) (Gadea and Lopez-Colome, 2001) (Fig. 5B), causing a manic phenotype, is inversely paralleled by a previous study. Helpless rats, an animal model for depression, displayed reduced expression of *Gat3* and showed depression-like behavior (Zink et al., 2009). Furthermore, alterations of glial function are considered to modify glutamate reuptake (Choudary et al., 2005), which may be the reason why we observed reduced glutamate levels in the NAc of our G*Per2* knock-out animals (Fig. 5I). Interestingly, however, we did not observe changes in expression of glutamate transporters *Eaat1* and *2* (Supplemental table 2). This is in contrast with our previous study investigating whole-body *Per2* mutant mice, which show decreased *Eaat1* levels (Spanagel et al., 2005). Hence, it appears that lack of *Per2* function in neurons and astrocytes leads to changes in *Eaat1* expression, illustrating the complex interplay between neurons and astrocytes in the regulation of glutamate signaling and mood-related behavior.

Deletion of *Bmal1* from astrocytes resulted in a change of GABA signaling accompanied by alteration in daily locomotor activity and cognitive functions (Barca-Mayo et al., 2017). Interestingly, *Gat1* as well as *Gat3* were altered in expression, comparable to our observations in the G*Per2* mice (Fig. 5A, B), although the time point of analysis was different between the two studies. However, this indicates that astrocytic GABA transporter activity appears to be important for a number of behaviors, including mood-related behaviors, which could be linked to excitatory neurotransmission (Boddum et al., 2016).

Overall, our results provide evidence for a specific role of glial *Per2* in mood-related behavior, accompanied by dysregulation of components of the glutamatergic and GABAergic signaling pathways.

## Materials and Methods

### Animals and housing

All mice were housed with food and water *ad libitum* in transparent plastic cages (267 mm long × 207 mm wide × 140 mm high; Techniplast Makrolon type 2 1264C001) with a stainless-steel wire lid (Techniplast 1264C116), kept in light-and soundproof ventilated chambers. All mice were entrained to a LD 12:12 cycle, and the time of day was expressed as zeitgeber time (ZT; ZT0 lights on, ZT12 lights off). Two- to four-month-old males and females were used for the experiments unless otherwise stated. Males were used for the FST, while females were used for the O-maze and mixed genders to monitor activity. Housing as well as experimental procedures were performed in accordance with the guidelines of the Schweizer Tierschutzgesetz and the Declaration of Helsinki. The state veterinarian of the Canton of Fribourg approved the protocols.

Conditional glial *Per2* knock-out animals were generated using *Gfap*-*Cre* mice (Jackson lab FVB-Tg(GFAP-cre)25Mes/J, stock no. 004600, created by the laboratory of A. Messing) that were cross-bred with our *Per2* floxed animals ((Chavan et al., 2016), European Mouse Mutant Archive (EMMA) strain ID EM: 10599, B6;129P2-Per2^tm1Ual^/Biat). The resulting *GPer2* line was back-crossed to the C57BL/6 strain. The *Per2*^*Brdm1*^ mutant mice (Zheng et al., 1999) and their wild-type littermate controls were on a mixed 129 and C57BL/6 background.

### Viruses used and their application

The following viruses, produced by the Viral Vector Facility of the Neuroscience Center Zurich, were used: v95-PHP.eB (ssAAV-PHP.eB/2-hGFAP-EGFP-WPRE-hGHp(A)) as the control virus and v232-PHP.eB (ssAAV-PHP.eB/2-hGFAP-EGFP_iCre-WPRE-hGHp(A)) as the virus for recombination, v344-9 (ssAAV-9/2-shortCAG-chI[1x(shm/rNS)]-EGFP-WPRE-SV40p(A)) and v25-PHP.eB (ssAAV-PHP.eB/2-CAG-EGFP_Cre-WPRE-SV40p(A)) as controls for BBB permeability determination, while stereotactic injections were also performed with constructs v95 and v232 in the AAV9 capsid. All viruses expressed the green fluorescent protein as a marker.

For intravenous delivery of viruses, mice were sedated and chemically restrained with an intraperitoneal injection of 40 mg per kg body mass ketamine and 0.15 mg per kg medetomidine in saline and placed on a heating pad, while hydrogel was applied to their eyes. We injected 10^11^ viral genomes per mouse in a total volume of 200 μl PBS via the lateral tail vein. The sedation was reversed with atipamezole in 5 times the dose of medetomidine.

Whole-brain fluorescent imaging was performed using the IVIS Lumina II (Caliper LifeSciences) and the accompanying Living Image software (version 4.2.0.14335). Settings: excitation 465nm, filter GFP, exposure time 1 s, light level low, binning low/small.

### Forced swim test (FST)

The Porsolt’s forced swim test has been described in detail elsewhere (Hampp et al., 2008; Porsolt et al., 1978). Briefly, the mice were placed into a cylinder filled with water, where they were left for 6 min. The first 2 min were discarded as adaptation period, while the following 4 minutes were scored. The test was performed for four consecutive days. The first day is regarded as adaptation and was discarded. The following three days were manually scored for immobility time – the amount of time a mouse passively floated during the 4 min test window. An average immobility time for each mouse was calculated and then these were pooled according to the assigned experimental group. In the case of animals injected with the PHP.eB virus, the forced swim test was performed 4 weeks post injections (Challis et al., 2019). All forced swim tests were performed at ZT 6, 6 hours after lights were switched on in a 12-hour light, 12-hour dark environment (Hampp et al., 2008). The water temperature was always adjusted to 26 +/-1 °C.

The FSTs were recorded from the side view. The automated forced swim test analysis was performed with a custom written program that evaluated the relative pixel difference in a rectangular area spanning the width of the swim tank and having the height equal to one sixth of the width, measured from the water surface towards the bottom of the tank. Each pixel in the test area was evaluated for changes between frames in red, green and blue on the decimal scale, the differences were averaged and divided by the test area, resulting in a graph that showed relative pixel change per area over time.

### O-maze

The anxiety-based O-maze test was performed on an elevated 5.5 cm wide circular runway with an outer diameter of 46 cm, which was divided into 4 sections, with 2 opposing closed sections. The time spent in the open sections was evaluated based on video recording. Entry into the open section was considered when the mouse entered it with all 4 paws. To avoid habituation to the test environment, a single test of 5 min was performed.

### Tissue isolation, gene expression analysis and neurotransmitter quantification

For both gene expression analysis and neurotransmitter quantification, fresh brain tissue was dissected and immediately submerged into liquid nitrogen. For the list of primers, please see Supplemental table 4. For gene expression analysis, RNA was isolated using the Macherey-Nagel RNA Plus kit, and reverse transcribed using the Invitrogen SuperScript II. qPCRs were performed using the RotorGene 6000, with the KAPA Probe or KAPA SYBR master mix reagent.

### Surgical procedures

The surgical procedures were performed as previously described (Martini et al., 2019). Briefly, mice were anesthetized with 80 mg per kg ketamine and 0.30 mg per kg medetomidine, while the anesthesia was reversed by atipamezole in 5 times the dose of medetomidine. Before surgical procedures, the depth of anesthesia was checked by an absence of a reflex when pinching the skin between the toes of the mouse 5-10 min after application of the anesthetic. In case of a reflex, a quarter of the initial dose of the anesthetic was additionally administered. For measurements of internal body temperature and general activity, a wireless biochip (VitalView system) was implanted into the abdominal cavity (Starrlife Sciences, VitalView Data Acquisition System, Instruction Manual, Software Version 5.1). Stereotactic injections were performed with a pulled glass pipette, which allowed injections of 2 × 200 nl of the virus bilaterally into the nucleus accumbens (NAc) with stereotactic coordinates of 1.60 anterior-posterior, +/-1.00 medio-lateral and 4.50 dorso-ventral. Surgical procedures were followed by administration of carprofen at 10 mg per kg.

### Immunohistochemistry

Brains were harvested after mice were cardiovascularly perfused with saline and 4% PFA, and the tissue was left in PFA over night, and then transferred to 30% sucrose in PBS for two days for cryoprotection. 40 µm sections were cut using a cryostat and kept at -20 °C in an antigen preservation solution (1% m/m polyvinyl pyrrolidone in a 1:1 mixture of PBS and ethylene glycol) until subsequent steps of washing, blocking with permeabilization, antigen retrieval and antibody staining (anti-PER2, Alpha Diagnostic, cat. PER21-A, 1:200 dilution; anti-GFAP, Abcam, cat. ab53554, 1:500 dilution; anti-NeuN, Merck Millipore, cat. mab377, 1:250 dilution). The procedures have been described by our laboratory in detail elsewhere (Brenna et al., 2019). Images were acquired using a Leica SP5 confocal microscope.

### Wheel-running experiments, general activity and internal body temperature measurements

Wheel-running experiments were performed using custom built cages, according to local legislation on animal experimentation, which had a stainless-steel wire running wheel with a diameter of 11.5 cm. On the axis, a system with a magnet closed a switch for each wheel revolution. The data was digitalized using an interface from Actimetrics and the activity was recorded and processed using the ClockLab software version 6.0.54.

This allowed acquisition of activity patterns in LD conditions, as well as monitoring of the subjective day length under constant conditions. For this, the mice were entrained to LD conditions and then released into total darkness for a minimum of 10 days. The first three days were discarded as assimilation, while the subsequent 7 days were evaluated for a shift of activity onset on each consecutive day in order to give a measure of the length of the subjective day or free-running period, as previously described (Jud et al., 2005; Martini et al., 2019).

Telemetrics was performed with the VitalView system, which is used for wireless measurements of general mouse activity, as well as internal body temperature, allowing monitoring of behavioural and physiological changes in free-running animals. For the biochip implantation, please see ‘Surgical procedures’. All the procedures have been described in detail elsewhere (Martini et al., 2019).

### Flow cytometry

At the desired time-point, brain tissue was harvested, dissected, stored in a test tube and submerged into cooled isopentane. The samples, still in the isopentane bath, were then stored at -80 °C (Rubio et al., 2016). For flow cytometry, the samples were diced with razor blades and digested under heat and agitation with the Neural Tissue Dissociation Kit (T) (130-093-231) from Miltenyi Biotech according to the manufacturer’s protocol. Single-cell suspensions were obtained by straining through a 70 µm mesh filter, after which the strained cells were washed twice in FACS buffer (PBS with 5% fetal calf serum and 5mM EDTA). Then, myelin debris were removed using the Myelin Removal Beads II kit (130-096-731) from Miltenyi Biotech. Myelin-depleted cell suspensions were incubated with an anti-CD16/CD32 Fc-blocking antibody (BD Biosciences) for 30 minutes at 4 °C to avoid Fc receptor binding and then washed once with FACS buffer. The cells were stained with fluorophore-conjugated antibodies and analyzed on a MACSQuant flow cytometer (Miltenyi Biotech) or sorted using the FACSAria Fusion cell sorter (BD Biosciences). Flow cytometry analysis was based on viable and single-cell gaiting strategies, as previously described (Schwarz et al., 2013). Antibodies targeted CD11b (fluorophore PECy7, ref. 552850, BD Pharmingen), CD90.2 (fluorophore PE, ref., 130-102-489, Miltenyi Biotech) and GLT1 (fluorophore ATTO 633, ref., AGC-022-FR, Alomone lab), and were used in a 1:100 dilution.

## Acknowledgments

We would like to acknowledge the invaluable advice from Prof. Jaclyn Schwarz from the University of Delaware on flow cytometry of brain tissue samples. S. Cattin and O. Coquoz from the Cell Analytics facility of the University of Fribourg are acknowledged for technical assistance and guidance for flow cytometry analysis and cell sorting experiments. We would also like to thank Dr. Jean-Charles Paterna from the Virus Vector Facility of the University of Zurich and ETH Zurich for providing valuable advice regarding the construction of viral vectors. We also thank A. Hayoz, S. Aebischer, and A. Sargsyan for technical support. AAV-pgk-Cre was a gift from Patrick Aebischer (Addgene viral prep # 24593-AAVrg).

## Notes

### Competing Interest Statement

The authors have declared no competing interest.

