## Supplemental Information for "Deletion of the Clock Gene *Period2 (Per2)* in Glial Cells Alters Mood-Related Behavior in Mice"

### Supplemental Figure Legends

#### Supplemental Figure 1

Swimograms of all PHP.eB-mediated glial KO animals (*Per2* $^{fl/fl}$ *Gfap-iCre-eGfp*) on the final day of the forced swim test (green) vs. PHP.eb injected controls (*Gfap-eGfp*)(blue) show a lack of longer stretches of immobility time in the KO group, with prolonged stretches of low activity in the control group.

**Supplemental Tables**

**Supplemental Table 1: Statistics of germline recombination in different families of G*Per2* mice.**

A simple determination of the percent of mice with germline recombination was based on the frequency of the deleted *Per2* allele in *Cre*-negative offspring. This frequency was individually evaluated for families where the *Cre* carrier was the male or female breeding partner, respectively. Only families where both alleles were floxed in the mating partners were analyzed.

**Supplemental Table 2: Gene expression analysis in the NAc, mPFC, AMY and HYP.**

Changes in gene expression in different brain regions. The relative change was calculated by dividing the relative mRNA in the G*Per2* by that in the control group, and the group comparison was made with the two-tailed Student's t-test.

**Supplemental Table 3: Glu, Gln and GABA quantification in the NAc, DS and mPFC.**

Freshly isolated and flash-frozen tissue was used for total (intra- and extracellular) quantification of neurotransmitter content at ZT6. The results are given in both nanomoles and nanograms per mg of isolated tissue in control animals (C) and G*Per2* animals (KO). The group comparison was made with the two-tailed Student's t-test.

**Supplemental Table 4: Primers for PCR.**

The TM probes were synthesized with a 6-fluorescin (FAM)group at the 5’-end and a black hole quencher 1 (BHQ1) group at the 3’-end. All other primers were unmodified.

| | | Progeny | | | | | | | | | | |
|---|---|---|---|---|---|---|---|---|---|---|---|---|
| | | *Cre +/-* | | | *Cre -/-* | | | | *Cre +/-* | *Cre -/-* | | |
| **Mating** | ***Cre* carrier** | **fl/fl** | **fl/Dfl** | **total** | **fl/fl** | **fl/Dfl** | **total** | **n** | **% Dfl** | **% Dfl** | **Difference** | **Notes** |
| 26 | female | 10 | 27 | 37 | 18 | 25 | 43 | 80 | 72.97 | 58.14 | 14.83 | |
| 29 | male | 1 | 20 | 21 | 22 | 0 | 22 | 43 | 95.24 | 0.00 | 95.24 | |
| 33 | male | 1 | 2 | 3 | 5 | 0 | 5 | 8 | 66.67 | 0.00 | 66.67 | |
| 36 | male | 3 | 15 | 18 | 24 | 0 | 24 | 42 | 83.33 | 0.00 | 83.33 | |
| 38 | male | 6 | 4 | 10 | 19 | 0 | 19 | 29 | 40.00 | 0.00 | 40.00 | |
| 41 | male | 1 | 2 | 3 | 6 | 0 | 6 | 9 | 66.67 | 0.00 | 66.67 | |
| 44 | male | 7 | 17 | 24 | 15 | 0 | 15 | 39 | 70.83 | 0.00 | 70.83 | |
| 45 | male | 35 | 14 | 49 | 41 | 0 | 41 | 90 | 28.57 | 0.00 | 28.57 | 1 discarded (inconclusive genotype). |
| | | | | | | M carrier | 132 | 260 | 64.47 | 0.00 | | |
| | | | | | | F carrier | 43 | 80 | 72.97 | 58.14 | | |

| | | nucleus accumbens | | | amygdala | | | medial prefrontal cortex | | | hypothalamus | | | |
|---|---|---|---|---|---|---|---|---|---|---|---|---|---|---|
| gene | full name | rel. Δ vs. WT * | p-value ** | N per group | rel. Δ vs. WT * | p-value ** | N per group | rel. Δ vs. WT * | p-value ** | N per group | rel. Δ vs. WT * | p-value ** | N per group | smallest p-value |
| Gat3 | GABA transporter 3 | 1.67200 | 0.00792 | 6 | 0.84251 | 0.65663 | 3 | 1.12610 | 0.42204 | 3 | 0.49669 | 0.15719 | 3 | 0.00792 |
| Eaat1 | excitatory amino acid transporter 1 | 1.04613 | 0.85813 | 6 | 0.95439 | 0.76588 | 3 | 1.02759 | 0.95246 | 3 | 0.99692 | 0.97981 | 3 | 0.76588 |
| Eaat2 | excitatory amino acid transporter 2 | 0.92958 | 0.66133 | 6 | 0.90787 | 0.57846 | 3 | 1.03055 | 0.91148 | 3 | 0.87722 | 0.12746 | 3 | 0.12746 |
| Maoa | monoamine oxidase A | 0.99121 | 0.95198 | 6 | 1.01672 | 0.93794 | 3 | 1.00434 | 0.98245 | 3 | 1.00145 | 0.99466 | 3 | 0.93794 |
| Maob | monoamine oxidase B | 0.94880 | 0.76553 | 6 | 0.91038 | 0.18386 | 3 | 1.08833 | 0.67899 | 3 | 1.12360 | 0.28016 | 3 | 0.18386 |
| Gabra1 | GABA receptor subunit alpha-1 | 0.97835 | 0.90355 | 6 | 0.95540 | 0.69682 | 3 | 0.92147 | 0.76567 | 3 | 0.92538 | 0.50802 | 3 | 0.50802 |
| Gabra2 | GABA receptor subunit alpha-2 | 1.14240 | 0.28715 | 6 | 1.51702 | 0.05586 | 3 | 1.52571 | 0.10686 | 6 | 1.45088 | 0.11052 | 3 | 0.05586 |
| Gat1 | GABA transporter 1 | 0.97738 | 0.94776 | 6 | 1.15281 | 0.58424 | 3 | 1.13179 | 0.51951 | 3 | 0.49651 | 0.02529 | 3 | 0.02529 |
| Aadc | aromatic L-amino acid decarboxylase | 1.47268 | 0.52475 | 3 | 0.93074 | 0.81351 | 3 | 0.88371 | 0.54718 | 3 | 0.72737 | 0.31008 | 3 | 0.31008 |
| Tph2 | tryptophan hydroxylase 2 | 1.90991 | 0.21618 | 3 | 0.80914 | 0.51223 | 3 | 0.87692 | 0.74469 | 3 | 0.99168 | 0.95876 | 3 | 0.21618 |
| Th | tyrosine hydroxylase | 1.01326 | 0.96115 | 6 | | | | | | | | | | 0.96115 |
| Comt | catechol-O-methyltransferase | 1.01111 | 0.91634 | 6 | | | | | | | | | | 0.91634 |
| Gls | glutaminase | 0.99761 | 0.99229 | 6 | | | | | | | | | | 0.99229 |
| Glns | glutamine synthetase / glutamate ammonia ligase | 1.07378 | 0.45718 | 6 | | | | | | | | | | 0.45718 |
| Drd1 | dopamine receptor D1 | 1.01636 | 0.89078 | 6 | | | | | | | | | | 0.89078 |
| Drd2 | dopamine receptor D2 | 1.05641 | 0.71733 | 6 | | | | | | | | | | 0.71733 |
| Drd3 | dopamine receptor D3 | 2.11310 | 0.00081 | 6 | | | | | | | | | | 0.00081 |
| Trank1 | tetratricopeptide repeat and ankyrin repeat containing 1 | 1.20527 | 0.40539 | 6 | | | | | | | | | | 0.40539 |
| Tac1 | tachykinin precursor 1 | 1.01403 | 0.89458 | 6 | | | | | | | | | | 0.89458 |
| Per2 | period 2 | 0.66225 | 0.00442 | 6 | | | | | | | | | | 0.00442 |
| | | | | | | | | | | | | | | 0.00000 |
| | | | | | | | | | | | | | | 0.00000 |
| | | | | | | | | | | | | | | 0.00000 |
| | | | | | | | | | | | | | | 0.00000 |
| | | | | | | | | | | | | | | 0.00000 |
| | | | | | | | | | | | | | | 0.00000 |
| | | | | | | | | | | | | | | 0.00000 |
| | | | | | | | | | | | | | | 0.00000 |

* expression in WT is 1, therefore a reduction is < 1 and an upregulation is > 1

** two-tailed student's t-test

| | | dorsal striatum | | | | | | nucleus accumbens | | | | | | medial prefrontal cortex | | | | | |
|---|---|---|---|---|---|---|---|---|---|---|---|---|---|---|---|---|---|---|---|
| neurotransmitter | full name | nmol/mg C | nmol/mg KO | ng/mg C | ng/mg KO | p-value ** | N per group | nmol/mg C | nmol/mg KO | ng/mg C | ng/mg KO | p-value ** | N per group | nmol/mg C | nmol/mg KO | ng/mg C | ng/mg KO | p-value ** | N per group |
| GABA | gamma-aminobutyric acid | 8.15295 | 7.33702 | 840.73239 | 756.59351 | 0.5880459 | 5 C, 3 KO | 5.07252 | 4.36163 | 523.07840 | 449.77147 | 0.658647 | 5 C, 5 KO | 7.95110 | 6.86236 | 819.91757 | 707.64669 | 0.942352 | 8 C, 7 KO |
| glu | glutamate | 8.95074 | 8.43552 | ########### | 1241.118 | 0.5197857 | 5 C, 3 KO | 9.25206 | 8.55005 | ########### | 1257.9682 | 0.0377162 | 5 C, 5 KO | 11.39612 | 11.03182 | ########### | 1623.1119 | 0.159267 | 10 C, 8 KO |
| gln | glutamine | 5.28297 | 5.67295 | 772.05306 | 829.0453 | 0.5706658 | 5 C, 3 KO | 5.70956 | 5.37422 | 834.39564 | 785.3882 | 0.3380539 | 5 C, 5 KO | 4.22981 | 4.27389 | 618.14510 | 624.5863 | 0.829856 | 10 C, 8 KO |

** two-tailed student's t-test

**List of Primers:**

***Aadc***

FW: CAT GAG AGC TTC TGC CCT TCG G
RV: GCA GGA TGT GGT CCC CAG TGT
TM: CGG GAC AAG GCA GCT GGC CTG ATT CCA

***Comt***

FW: GTG GCT ACT CAG CCG TGC GA
RV: GCT GGG TGA TGG CAG CGT AGT
TM: TGG CCC GCC TGC TGC CAC CT

***Ddc***

FW: AGA AGA ACT GGT GTG AGG AGC AGT
RV: TGC CTG CAG CTG GCG GAT AA
TM: TGG TGG CCC TAC TGG CTG CTC GGA

***Drd1***

FW: AGG AGA GGG CGC AGG GTT G
RV: GCC CCT GGT GCC ACA TCT CT
TM: CGG AGT CGG GGA GCG TGG TCT CCC

***Drd2***

FW: GAT GCG GCG GGA GCT GGA A
RV: TGG GTG GCA CGG CTC TTC AA
TM: TCT CTG GCC CCG GGC GCC CT

***Drd3***

FW: CCT GGC TTC CCT CAG CAG TCT
RV: GCT CCA TTT GTC CCG TGG CAT CT
TM: TGT CTG CGG CTG CAT CCC ATT CGG CA

***Eaat1***

FW: GAT GCT GGT CTT GCC CCT GAT
RV: ACA GCG CGC ATC CCC ATC TT
TM: CCA GTC TCG TCA CAG GAA TGG CGG CCC

***Eaat2***

FW: ACA GGG TTG TCA GGC CTG GAT
RV: CAG CAC GGC GGC AAT GAT GG
TM: AGC CAG CGG CCG CCT AGG CA

***Gabra1***

FW: GTC TGG AGC GAT CCG GTG C
RV: TGA GGG TCC AGG CCC AAA GA
TM: CCC GAG CTG TGC AAG CCC GTG ATG A

***Gabra2***

FW: TGC AAT GTA TGG TCT CTG CTG CTT GT
RV: AGC CTC ATC TTC TTG GAT GTT AGC CA
TM: TGG TGT GGG ACC CAG TCA GGT TGG TGC

***Gat1***

FW: CGT GGA ACA CTG ACC GCT GCT
RV: GGT GCA TGT TGC GCT CCC AGA
TM: CCA CCA ACA TGA CCA GCG CCG TGG TGG

***Gat3***

FW: CTG GGA GAG GCG AGT CCT GA
RV: GCA GGA GGC ACA GGA CCA GTT
TM: CGG ATG GCA TCC AGC ACC TGG GGT CC

***Glns***

FW: TGG ACC CCA AGG CCC GTA T
RV: ACA AGC AGG CCC GGT AGT GA
TM: AGG CCT TGT CTG CTC CCA CAC CGC A

***Gls***

FW: TGT CTG CCC TCC GAA GGT TTG C
RV: ACC CTC TGC TGC TGC GAC AT
TM: CTG TCA GCC ATG GAC ATG GAG CAG CGG G

***iCre***

FW: GGG TTA CCA AGC TGG TGG AG
RV: GGC AGC CAC ACC ATT CTT TC

***MaoA***

FW: GGT ATG TGA GGC AGT GTG GAG GT
RV: CAC TTA TTT GGC CAG AGC CAC CT
TM: CAG TCA CCA ATG GCG GCC AGG AAC GGA

***MaoB***

FW: TGG AGC GGC TAC ATG GAG GG
RV: TCT GGA ATC TTC CCA ATG GCA TGA AGA

TM: TGG AGG CTG GGG AGA GAG CAG CCA

***Per2***

FW: TCC ACA GCT ACA CCA CCC CTT A
RV: TTT CTC CTC CAT GCA CTC CTG A
TM: CCG CTG CAC ACA CTC CAG GGC G

***Tac1***

FW: AGA GCA AAG AGC GCC CAG CA
RV: CGC CAC GGC CAC GAG GAT TT
TM: CCT GCG GAG CAT CCC CGC GG

***Th***

FW: CCT GGA CCA TCC GGG CTT CT
RV: GGG GAA TTG GCT CAC CCT GCT T
TM: CCA GGC GTA TCG CCA GCG CCG G

***Tph1***

FW: ACT GCG ACA TCA GCC GAG AAC A
RV: TTC GCA GTG AGC TGA TCG GG
TM: ACG CCA CCG TCC TCT CGG TGG ACT C

***Tph2***

FW: ACC CAG TAC GTG CGG CAT GG
RV: GCA GTG GCA CGT GTC CCA AGA
TM: CCG ACC CCC TCT ACA CCC CGG AAC C

***Trank1***

FW: GGT GTC ACC TGC GCG CAT CC
RV: GCC AGT TCC CGA GGA GGA GT
TM: CGG CGG CGA GTC CCG GCC AT

***Tspo***

FW: GGT CAG CTG GCT CTG AAC TG
RV: CAG TCG CCA CCC CAC TGA CA
TM: TGC CCG GCA GAT GGG CTG GGC

The TM probes were synthesized with a 6-fluorescin (FAM)group at the 5′-end and a black hole quencher 1 (BHQ1) group at the 3′-end. All other primers were unmodified.

| Resource (strain, reagent etc.) | Designation | Source | Referrence or identifier | Additional information |
|---|---|---|---|---|
| M. musculus, Per2 mutant | Per2Brdm1 | Jackson Laboratory | Stock #: 003819 | PMID: 10408444 |
| M. musculus, Per2 floxed | Per2tm1Ual/Biat | opean mouse mutant arch | Strain ID EM: 10599 | PMID: 26838474 |
| M. musculus, Gfap-Cre | FVB-Tg(Gfap-Cre)25Mes/J | Jackson Laboratory | Stock #: 004600 | PMID: 11668683 |
| AAV-PHP.eB (Gfap-eGfp) | v95-PHP.eB | VVF Zurich | v95-PHP.eB | |
| AAV-PHP.eB (Gfap-iCre-eGfp) | v232-PHP.eB | VVF Zurich | v232-PHP.eB | |
| AAV9 (CAG-eGfp-Cre) | v344-9 | VVF Zurich | v344-9 | |
| AAV-PHP.eB (CAG-eGfp-Cre) | v25-PHP.eB | VVF Zurich | v25-PHP.eB | |
| AAV9 (Gfap-eGfp) | v95-9 | VVF Zurich | v95-9 | |
| AAV9 (Gfap-iCre-eGfp) | v232-9 | VVF Zurich | v232-9 | |
| antibody | anti-PER2 | Alpha Diagnostic | PER21-A | dilution 1 : 200 (IF) |
| antibody | anti-GFAP | Abcam | ab53554 | dilution 1 : 500 (IF) |
| antibody | anti-NeuN | Merck Millipore | MAB377 | dilution 1 : 250 (IF) |
| commercial kit | SuperScript II Reverse Transcriptase 10,000 units | Thermo Fischer | 18064022 | |
| commercial kit | Qiagen RNeasy Micro | Qiagen | 74004 | |
| commercial kit | Macherey-Nagel NucleoSpin RNA Plus | Macherey-Nagel | 741984.5 | |
| software for wheel-running | ClockLab Analysis | Actimetrics | 6.0.54 | |
| software for image export | Fiji | freeware | version 2.0.0-rc-69/1.52p | |
| software for statistical analysis | R Studio | freeware | version 1.2.1335 | |
| software for image acquisition | Leica application Suite | Leica | version 2.7.3.9723 | |
| commercial kit | Neural Tissue Dissociation Kit | Miltenyi Biotech | 130-093-231 | |
| commercial kit | Myelin Removal Beads II | Miltenyi Biotech | 130-096-731 | |
| antibody | anti-CD16/CD32 Fc-blocking antibody | BD Biosciences | 553141 | |
| antibody | anti-CD11b, fluorophore PECy7 | BD Biosciences | 552850 | dilution 1 : 100 (FC) |
| antibody | anti-CD90.2, fluorophore PE | Miltenyi Biotech | 130-102-489 | dilution 1 : 100 (FC) |
| antibody | anti-GLT1, fluorophore ATTO 633 | Alomone lab | AGC-022-FR | dilution 1 : 100 (FC) |
